# Evaluation of microbiome enrichment and host DNA depletion in human vaginal samples using Oxford Nanopore’s adaptive sequencing

**DOI:** 10.1101/2021.09.15.460450

**Authors:** Mike Marquet, Janine Zöllkau, Jana Pastuschek, Adrian Viehweger, Ekkehard Schleußner, Oliwia Makarewicz, Mathias W. Pletz, Ralf Ehricht, Christian Brandt

## Abstract

Metagenomic sequencing is promising for clinical applications to study microbial composition concerning disease or patient outcomes. Alterations of the vaginal microbiome are associated with adverse pregnancy outcomes, like preterm premature rupture of membranes and preterm birth. Methodologically these samples often have to deal with low relative amounts of prokaryotic DNA and high amounts of host DNA (> 90%), decreasing the overall microbial resolution. Nanopore’s adaptive sampling method offers selective DNA depletion or target enrichment to directly reject or accept DNA molecules during sequencing without specialized sample preparation. Here, we demonstrate how selective ‘human host depletion’ resulted in a 1.70 fold (± 0.27 fold) increase in total sequencing depth, providing higher taxonomic profiling sensitivity. At the same time, the microbial composition remains consistent with the control experiments. The complete removal of all human host sequences is not yet possible and should be considered as an ethical approval statement might still be necessary. Adaptive sampling increased microbial sequencing yield in all 15 sequenced clinical routine vaginal samples, making it a valuable tool for clinical surveillance and medical-based research, which can be used in addition to other host depletion methods before sequencing.

## 1 Introduction

Long read sequencing technologies, such as nanopore sequencing, allows for fast, real-time, and culture-free metagenomic sequencing [1]. However, in samples containing relatively high proportions of host DNA (> 90%) like saliva, throat, buccal mucosa, and vaginal samples, the detection of low abundant species is expected to be impaired [2]. Host DNA depletion prior to sequencing by selective lysis of host and microbial cells or selective removal of CpG-methylated host DNA are complex wet-lab procedures [3]. For instance, host DNA depletion *via* saponin improves the sensitivity of pathogen detection after sequencing [4,5].

Oxford nanopore technologies (ONT) recently (November 2020) introduced target enrichment or depletion of unwanted DNA molecules (e.g., human DNA) directly during sequencing. While a DNA molecule is sequenced in the nanopore, the data is already compared live with references to decide whether the DNA molecule should be sequenced further (accepted or no decision yet) or removed directly from the pore (rejected). Each pore is individually addressable and can reverse the voltage on its pore to reject DNA molecules and sequence another one instead, increasing the sequencing capacity for molecules of interest [6]. The main advantage is that this depletion or enrichment method can be combined in addition to wet-lab depletion or enrichment methods and does not prolong the overall sequencing run time.

The host depletion by adaptive sequencing and thus enriching microbial organisms is likely to be significant for clinical samples, particularly if high levels of human DNA is expected (e.g., vaginal microbiome samples). The vaginal microbiome is characterized by low-alpha diversity (number of taxonomic groups) with a high relative abundance of *Lactobacillus* species. *Lactobacilli* promote vaginal and reproductive health producing specific metabolites (e.g., lactic acid, hydrogen peroxide, or bacteriocins) that inhibit colonization of the vagial micorenvironment by harmful microbiota [7]. Disruption or imbalance of the composition of vaginal microbiome during pregnancy (e.g., by antibiotic treatment) can result in complications such as preterm premature rupture of membranes (PPROM), cervical insufficiency, pregnancy loss, or preterm birth [8]. The latter is associated with early-onset neonatal sepsis (EONS) and risk of neonatal morbidity, mortality and may lead to long-term complications and deficits for the newborn [9–11]. Increased organism diversity and relative abundance of organisms like Group B streptococci, *Escherichia coli, Pseudomonas aeruginosa*, or *Ureaplasma parvum* and the depletion of certain *Lactobacilli* are indicators for bacterial vaginosis [7,12]. However, their identification *via* conventional culture-based, microbiological diagnostic techniques suffers from long reporting delays and low sensitivity and specificity [13,14]. On the other hand, pathogen identification based on 16S ribosomal RNA gene sequencing gives insights into the metagenomic composition with a lower resolution than metagenomics and suffers from various inherent biases to interpret abundance data properly: e.g., primer mismatches, different gene copy numbers, recombinations, sequence- and primer-dependent polymerase efficiency, or choice of hypervariable regions [15–18]. Nanopore sequencing has become a widely used method for metagenomic sequencing with similar taxonomic classification performance as short read sequencer (e.g., Illumina) [19–21]. The lower raw read accuracy of 97.8% is compensated *via* the higher information richness due to the longer reads. In the present study, we evaluated the overall performance of adaptive sampling to enrich bacterial reads and deplete human DNA in 15 human vaginal samples. The aim was to evaluate the strength and limitations of this technology and its potential metagenomics-based clinical diagnostic routines in culture-free metagenomic samples.

## 2 Results & Discussion

Vaginal samples from swabs mostly yield small amounts of DNA (< 40 ng/μl) for library preparation and subsequent sequencing, making PCR amplification often mandatory, which can introduce amplification bias and alter the microbial composition.

### 2.1 Influence of nonspecific amplification-based library preparation for the determination of microbial communities

The nanopore amplification-based library preparation kit (RPB004) uses transposase-mediated cleaving of DNA molecules to attach the primer binding sites for PCR amplification, which should reduce PCR amplification bias. We initially assessed this bias by determining the microbial composition of the ZymoBIOMICS Microbial Community Standard [22] (control) by sequencing using the RPB004 PCR-based library preparation kit and compared the abundance of the different species to the native PCR-free library preparation kit (LSK109). Both sequencing approaches were performed in triplicates to address experimental variations. The reads were mapped against the microbial genomes *via* minimap2 v.2.19 [23], counted *via* samtools depth v1.11 [24] (bases sequenced per organism), and summarised *via* ggplot2 (Figure 1).

**Figure 1:**
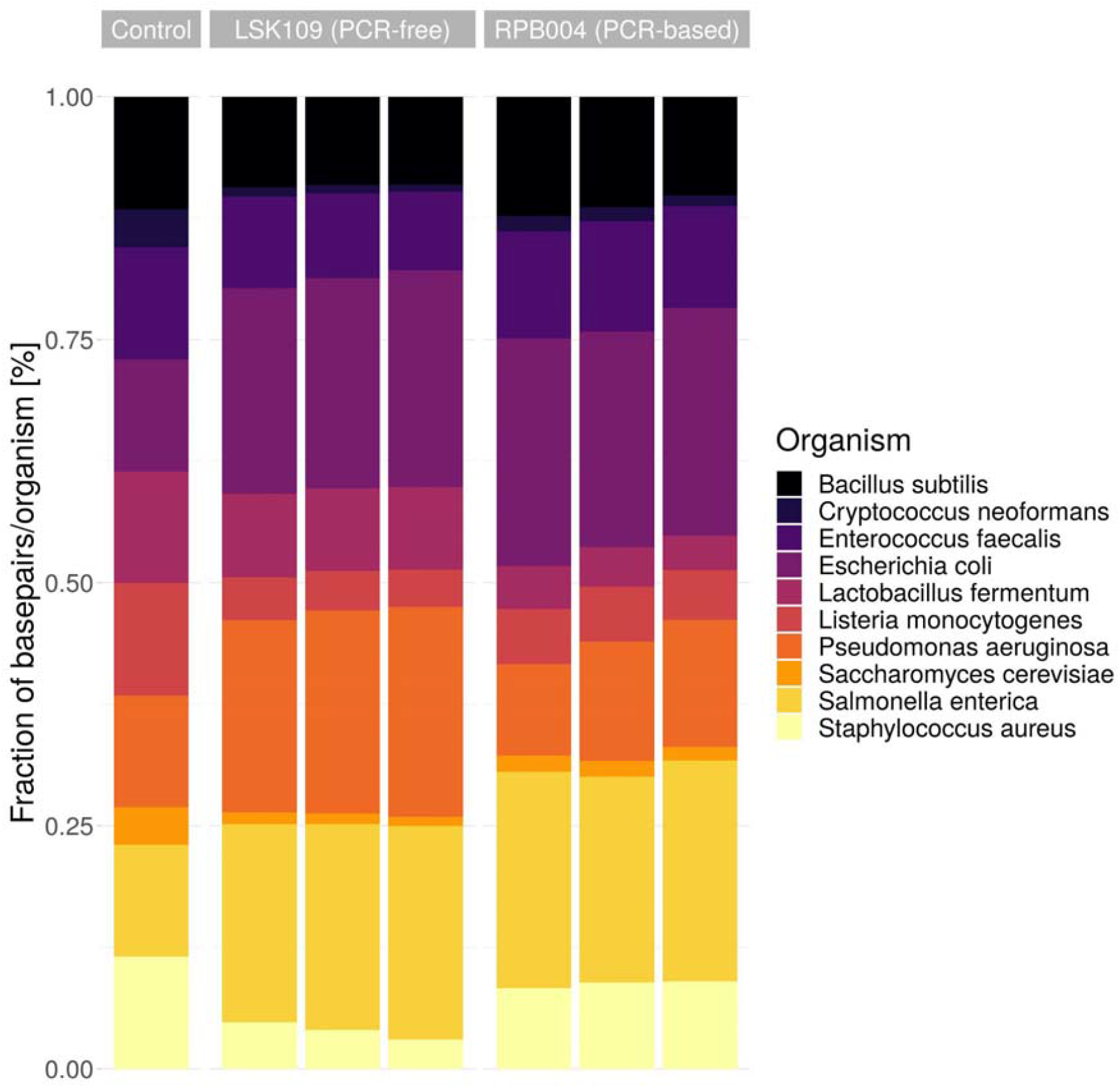
Abundance of the ten sequenced organisms of the ZymoBIOMICS Microbial Community Standard for the native PCR-free library preparation (LSK109) and the nanopore amplification-based library preparation (RPB004). The expected fraction for the microbial standard is shown on the left (control). The fraction of sequenced bases was determined by mapping the sequenced reads against the ten organisms via minimap2.

Both sequencing kits detected all ten organisms of the ZymoBIOMICS Microbial Community Standard and the results of the sample’s replicates exhibited only negligible deviation.

Compared to the expected abundance in the control, Gram-positive bacteria and yeast were underrepresented in the sequencing data obtained by both library preparation methods (amplification free: average 0.60 fold, min 0.34 fold, max 0.79 fold; amplification-based: average 0.70 fold, min 0.35 fold, max 0.97 fold), while Gram-negative bacteria were overrepresented (amplification free: average 1.84 fold, min 1.80 fold, max 1.88 fold; amplification-based: average 1.64 fold, min: 1.01 fold, max: 1.99 fold). The amplification-based library preparation approach shows a considerable difference to the PCR-free library preparation method for *Pseudomonas aeruginosa* (0.55 fold of PCR-based), *Lactobacillus fermentum* (0.47 fold of PCR-based), *Staphylococcus aureus* (2.21 fold of PCR-based), and *Cryptococcus neoformans* (1.57 fold of PCR-based). Six organisms show minor differences to the PCR-free library preparation *(Bacillus subtilis, Enterococcus faecalis, Escherichia coli, Listeria monocytogenes, Saccharomyces cerevisiae, Salmonella enterica*).

We expected the PCR-free library preparation approach to represent the microbial community standard more accurately since it was previously validated by other groups [25]. However, it showed clear variation in abundances compared to the control, which might be attributed to the different cell disruption device used in this work. We did not observe a significant advantage or disadvantage in choosing the amplification-based library preparation method over the PCR-free library preparation method to assess the microbial composition as both similarly overrepresent Gram-negative bacteria (Figure 1), but interestingly the PCR amplification-based library preparation seemed to represent the control slightly better than the PCR-free library preparation.

### 2.2 Adaptive sampling for metagenomes: Enrichment or depletion?

ONT’s adaptive sampling method enables it to either enrich or deplete DNA. To evaluate if microbial target enrichment or host depletion is more suitable for human vaginal metagenome sequencing (providing more microbial sequencing data), we compared both methods against a control experiment without adaptive sampling.

We sequenced a human vaginal metagenome (87.93% human host contamination) from a pregnant woman to derive species information for the enrichment process first. In a second step, we performed a depletion experiment using a human genome as reference (GCF_000001405.39). Finally, we performed an enrichment experiment using nine bacterial genomes downloaded from NCBI as reference based on the most abundant identified species from the first control sequencing experiment (see method section: Nanopore sequencing).

Each read passing the nanopore during adaptive sampling was mapped against a single or multiple reference genome(s) (e.g., human reference genome or multiple bacterial genomes) while sequencing. The mapping occurred in intervals of several bases, and three types of decisions were made: 1) ‘no_decision’ - the read has been continued and mapped against the reference(s) after several bases again (‘no decision’), 2) ‘stop_receiving’ - the read was accepted and fully sequenced (‘accepted’), 3) ‘unblock’ - the sequencing was immediately stopped and the read was rejected by reversing of the voltage (‘rejected’). The base pairs required until a decision has been made were summarised in Figure 2 B for all reads. For both methods, read rejections occurred within approx. 400 to 800 bp. Accepting reads started at approx. 400 bp or 4000 bp for the enrichment or depletion protocols, respectively. More generally, read lengths of at least 400 bp were required for both adaptive sampling methods to start the individual reads’ decision-making process.

**Figure 2:**
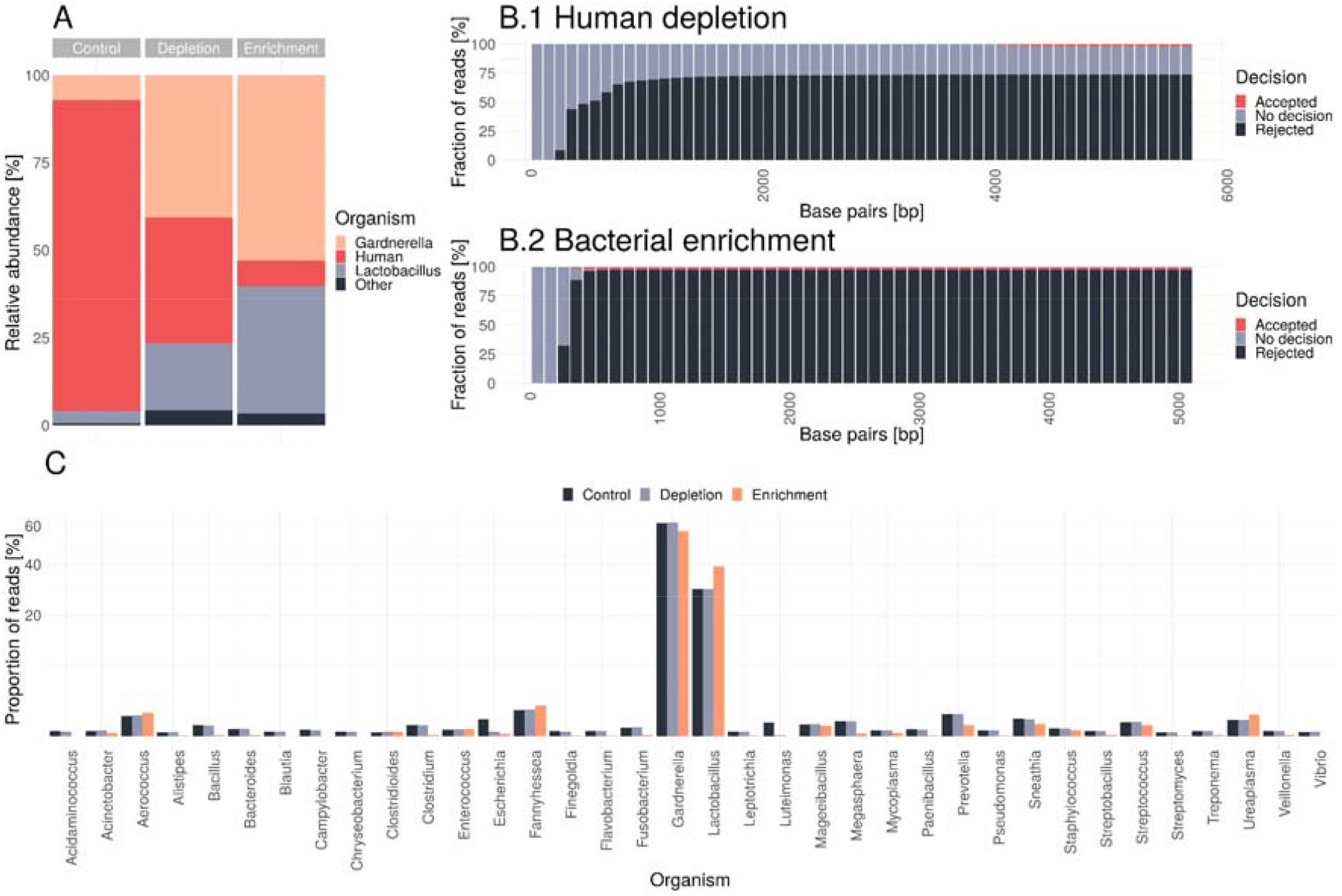
(**A**) Proportions of sequenced human, Gardnerella and Lactobacillus reads for the control, depletion and enrichment sequencing experiments using the ‘accepted’ and ‘no decision’ fractions for the adaptive sampling experiments only. Reads were taxonomically classified with centrifuge v1.0.4. (**B**) Base pairs required until a decision has been made in the depletion (**B.1**) and enrichment (**B.2**) experiment for all sequenced reads. (**C**) Proportion of sequenced genera for each of the experiments.

The enrichment experiment yielded a higher total reads’ number (5.67 million, 1.50 fold more than depletion, of which 5.44 million were rejected reads), followed by the depletion experiment (3.79 million reads, 1.39 fold more than the control, of which 3.07 million reads were rejected). The control yielded 2.73 million reads. One should note that experimental variations affect the total sequencing performance, but the yield increase *via* depletion of human sequences was further validated (see 2.3).

Due to short read lengths, which result from the high rejection rate and the fast decision process (Figure 2 B.2), the enrichment experiment yielded the least amount of total bases and microbial bases while ‘human depletion’ yields the most microbial bases (Table 1). Without adaptive sampling, the proportion of sequenced human reads was unsurprisingly highest (87.93%) but could be strongly reduced by the depletion approach to 34.73% and by the bacterial enrichment down to 8.29% (Figure 2 A). The ‘human depletion’ method rejected almost 81.01% of all reads, which was lower than the total abundance of human DNA in the control experiment, suggesting that the chosen human genome might be insufficient for a complete depletion of all human reads or the adaptive sampling process itself is prone to error. The bacterial enrichment method rejected 95.93% of all reads, which indicates that some bacterial reads were also rejected. We identified 5.48% of *Gardnerella* reads, 2.41% of *Lactobacillus* reads, and 2.20% of other microbial reads in the ‘rejected’ fraction of the bacterial enrichment experiment. Simultaneously, the proportions of essential vaginal microorganisms, like *Lactobacillus* and *Gardnerella*, could be increased by both methods but higher by the enrichment protocol.

**Table 1:** Summary of sequencing yield generated for the control, depletion and enrichment experiment using the PCR-based (RPB004) library preparation kit with a median read length of 2500 bp due to the amplification step. The calculated microbial bases are based on the ‘accepted’ & ‘no_decision’ fractions.

| Experiment | Total bases<br>[Gigabases] | Total reads<br>[million] | Rejected reads<br>[million] | Rejected<br>reads | microbial bases<br>[Gigabases] |
| --- | --- | --- | --- | --- | --- |
| Control | 9.91 | 2.73 | none | 0 % | 1.06 |
| Depletion | 5.2 | 3.79 | 3.07 | 81.01% | 1.43 |
| Enrichment | 4.26 | 5.67 | 5.44 | 95.93% | 0.77 |

We compared the proportions of bacterial genera of the experiments identified from the reads of the ‘accepted’ and ‘no decision’ category to validate whether the overall microbial composition was retained despite adaptive sampling (Figure 2 C). The human depletion method clearly showed very similar proportions to the control for 34 of 36 microorganisms except for *Escherichia* and *Luteimonas*. The difference in the proportions of these two organisms could be attributed to experimental variations, especially since their frequency in the control experiment was only 0.05% (*Escherichia*) and 0.03% (*Luteimonas*). Conversely, the enrichment method shows significant differences in most genera, including important vaginal microorganisms like *Gardnerella, Lactobacillus*, and *Ureaplasma*.

Therefore, we assume that an enrichment approach might be unsuitable for investigating the microbial composition between metagenomic samples if not all species can be reliably provided as target sequences during the enrichment. On the other hand, the ‘human depletion’ experiments maintained a comparable microorganism composition as the control experiments and considerably (53.20%) reduced the number of human reads, making it a robust choice for clinical metagenomic samples with high amounts of human host DNA.

### 2.3 Performance of human host depletion *via* adaptive sampling in human vaginal metagenomic samples

We collected 15 vaginal samples of pregnant women (see method section: Sample selection) with high proportions of host DNA (> 90%).

First, each of the 15 samples were sequenced without adaptive sampling serving as a control experiment and ground truth of their metagenomic composition to track possible changes introduced via adaptive sampling. We then sequenced the same isolated DNA from the control experiments while using adaptive sampling (human DNA depletion) and compared the overall sequencing performance to the previously sequenced controls (Figure 3). Additionally, a negative control of a swab without patient material following the same sample gathering and sequencing approach yielded 126 reads (ranging from 10 bp to 4000 bp) but none of the reads were classifiable and might be attributed to some PCR-primer and sequencing adapter constructs.

**Figure 3:**
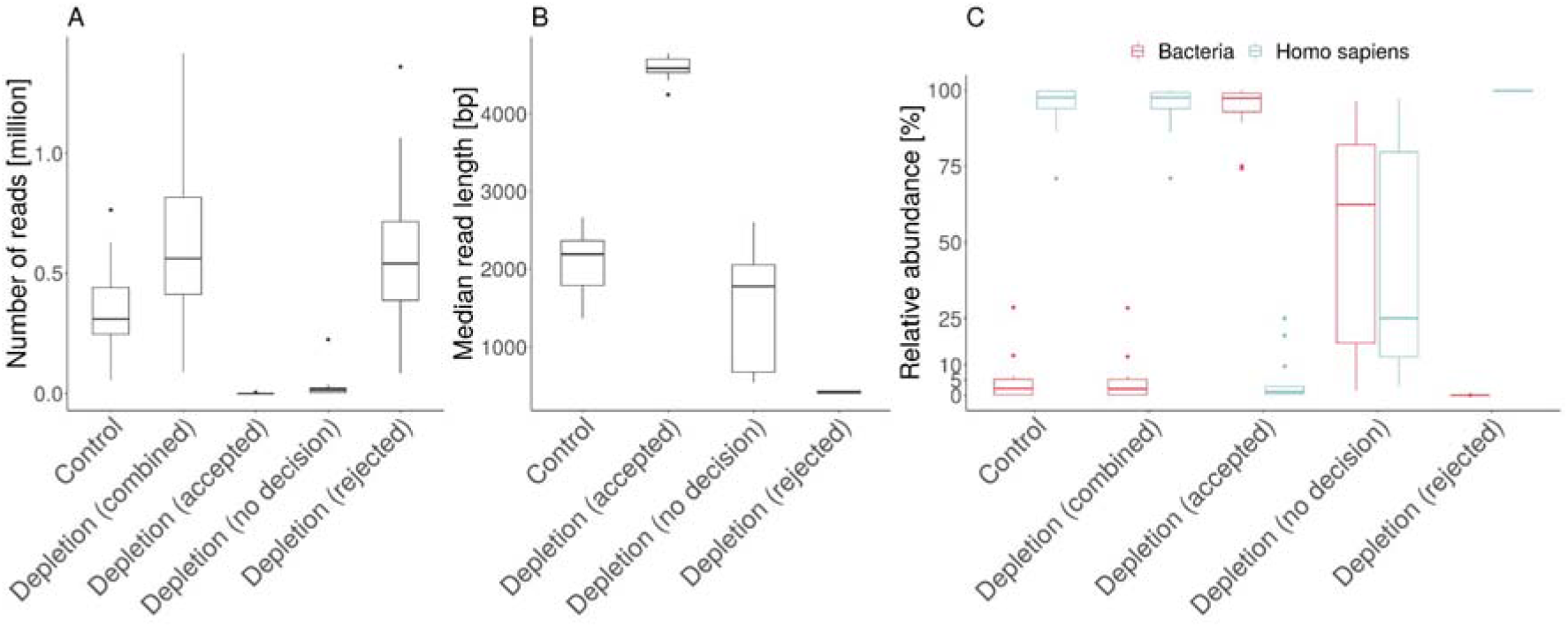
Overall performance of human depletion experiments compared to control experiments for 15 vaginal metagenomic samples. (**A**) Total number of sequenced reads for depletion and control experiments. Depletion experiments were additionally split into the three decision categories: ‘rejected’, ‘no decision’, and ‘accepted’. (**B**) Median read length distribution of the three depletion decision categories compared to the control experiments. (**C**) Human (blue) and bacterial (red) proportions for each sample of the control and depletion experiments. Depletion experiments were additionally split into the three decision categories: ‘rejected’, ‘no decision’, and ‘accepted’.

All reads were taxonomically classified *via* centrifuge v1.0.4 [26] to investigate their taxonomic composition (centrifuge database: Human-Virus-Bacteria-Archaea 01.2021) [27].

On average, depletion experiments yielded 1.71 fold (± 0.27 fold) more reads (including rejected reads) than the corresponding control experiments (Figure 3 A). This corresponds to a yield of ∼ 1.7 flow cells from a standard Nanopore sequencing experiment, with the only difference being that unwanted DNA molecules (human) were only partially sequenced. Reads of the categories ‘no decision’ (average: 5.38%, ± 7.24%) and ‘accepted’ (average: 0.23%, ± 0.35%) contributed a small overall proportion of all sequenced reads due to the high amount of human DNA. On average, adaptive sampling categorized 92.05% (± 7.42%) of reads as ‘rejected’. ‘Accepted’ reads were comparably long (Figure 3 C) with a median read length of ∼4000 bp, which is in line with previous results (Figure 2 B).

Samples of the control experiments contained 97.59% (± 7.63%) human reads, while bacteria reads made up to 2.22% (± 7.49%) (Figure 3 C). Without splitting into the three read categories, the depletion runs showed similar proportions of species like the control. The ‘rejected’ fraction of the depletion experiments contained 99.80% (± 0.09%) human reads and 0.02% bacterial reads (± 0.02%), indicating a very selective depletion process. Reads of the category ‘accepted’ contained low amounts of human reads 1.23% (± 8.13%) and 97.42% (± 8.38%) bacterial reads. Three samples contained 4, 20, and 92 reads only within the ‘accepted’-fractions and were excluded in the previous calculation. The ‘no decision’-fractions contained 62.37% bacterial reads (± 34.76%), but also 25.06% human reads (± 36.19%).

In summary, most of the not ‘rejected’ reads were placed into the ‘no decision’ category as the ‘accepted’ decision was rarely made. The ‘no decision’ category also included many human reads and most of the bacterial reads. Combining the ‘accepted’ and ‘no decision’ fractions of the bacterial reads generally yielded more microbial reads compared to the control experiment underlining the capabilities of adaptive sampling to increase the sequencing depth (Figure 3 C). Furthermore, the decision made for the ‘rejected’ category was remarkably accurate as it contained almost exclusively human reads. However, those human reads were still identified in the ‘accepted’ and ‘no decision’ fractions as observed in previous experiments. Thus, to reliably remove all human reads during sequencing seems rather elusive.

### 2.4 Depletion does not alter the species distribution in samples

Adaptive sampling selectively depletes human reads while simultaneously enriching microbial reads due to the increased sequencing depth but may alter species representation and metagenomic composition. We, therefore, compared the taxonomically classified reads of the 15 vaginal metagenomic samples in detail, to compare the individual abundance between control and the host depletion experiments (Figure 4 and Supplementary Figure S1 & S2).

**Figure 4:**
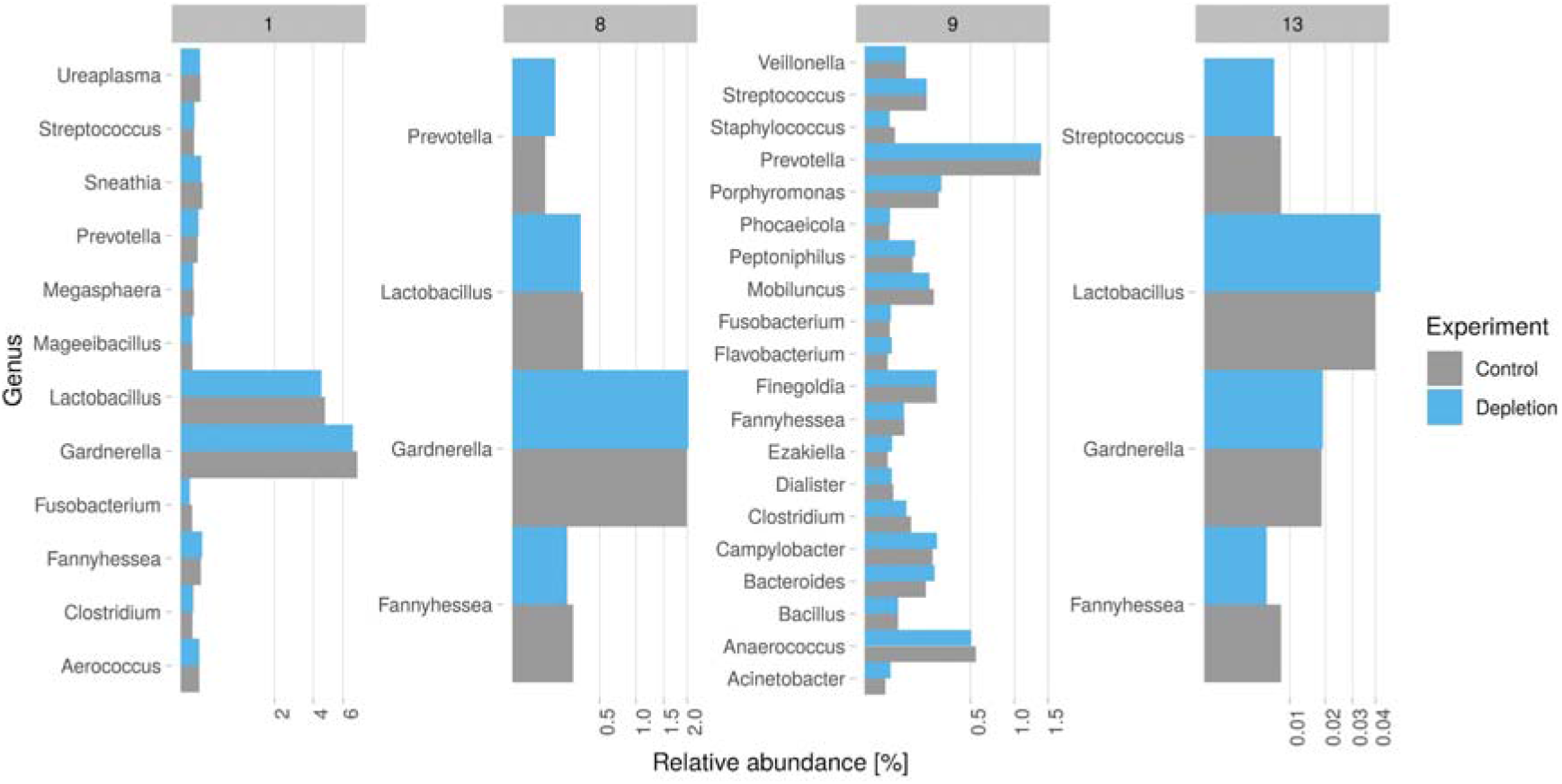
Comparison of the relative abundance of bacterial genera for four of the 15 vaginal samples (control in grey, depletion in blue). The full overview is shown in Supplementary Figure S1 for genus and Supplementary Figure S2 for species classifications.

The proportion of each genus was calculated in relation to the total amount of reads generated for each sample, including ‘rejected’ reads for the depletion experiment. We only included genera with at least 30 reads to avoid over-interpreting uncertain taxonomic classifications. This resulted on average in five bacterial genera (min 1, max 20) and in six bacterial species (min 1, max 26; Supplementary Figure S2), which correspond well with the expected vaginal microbiome [28]. Across all 15 samples, the bacterial proportion varied from the corresponding control experiments on average by 1.03 fold, indicating a similar representation of genus abundance levels by the depletion experiment (Figure 4). Low abundance genera with reads counts between >30 and <100 showed higher discrepancies (± 0.23 fold). Higher read counts showed higher reliability (e.g. >500 <1000 reads (± 0.03 fold) and >1000 reads (± 0.04 fold). This higher discrepancy in genera with a small number of reads is expected, as the experimental variability’s influence is more prominent. We did not detect any organisms that were found solely by only one method using the applied cutoff. Overall both experiment groups performed highly similarly in relation to the detected genus abundance levels between the control and depletion experiments.

## 3 Conclusion

The enrichment of targets in human cells [29–33] or species in mock communities [30,34] or fecal samples of lions [35] via adaptive sampling was previously demonstrated. In the present study, we used clinical metagenomic samples obtained from vaginal swabs of pregnant women to evaluate the performance to deplete the high content of human DNA that heavily impairs downstream microbiome analyses. Our results demonstrated that ONT’s unique adaptive sequencing feature has reliably increased the overall sequencing depth of bacterial sequences in clinical metagenomic samples *via* ‘human depletion’ without changing the microbial composition during sequencing. However, the enrichment experiment showed significantly higher ‘human depletion’, but changed the overall identified bacterial composition by also depleting other microbial sequences, illustrating that the enrichment method may be poorly suited for some microbiome studies.

Currently, to increase the sequencing depth for metagenomic samples with low microbial material, several sequencing runs per sample or sequencers with higher throughput (e.g., PromethION in case of ONT) are necessary, besides wet laboratory methods to deplete the host DNA. ONT’s adaptive sampling method demonstrated in our work a 1.7 fold increase in sequencing depth for samples with high human DNA contamination, which increases sensitivity of taxonomic profiling by providing more sequencing data [2]. Moreover, adaptive sampling can be used in addition to wet lab procedures to increase sensitivity further [36]. This makes molecular monitoring of human reservoirs with low microbial concentrations and high host DNA loads (e.g., nasal swabs, sputum, or skin swabs) more feasible. In this manner, the adaptive sampling improved the detected organisms beneficial to the vaginal microbiota, such as *Lactobacillus*, and pathogenic microbiota, such as *Streptococcus, Gardnerella*, or *Ureaplasma*, potentially harmful in pregnancy. Therapeutic measures can be derived based on the presence or ratio of certain species improving vaginal microecological diagnostics in the future, which enables clinically relevant insights into eubiosis or dysbiosis during pregnancy. On a side note, due to the increased sequencing depth *via* adaptive sampling, more samples can be barcoded and sequenced simultaneously, reducing the total cost per sample in a diagnostic laboratory.

However, human DNA could not be completely removed, so that the raw data always contained human sequences, which poses an ethical problem. Although the ONT adaptive sequencing could still be used for diagnostic purposes, patient consent is required for scientific purposes or data upload to public repositories like the National Center for Biotechnology Innovation (NCBI) or the European Nucleotide Archive (ENA). In turn, removing human sequences *by* different bioinformatics approaches poses hurdles for institutions and hospitals without adequate bioinformatics support. However, the research field of nanopore sequencing is evolving rapidly and dynamically, and ONT may address current limitations in the foreseeable future. In addition, continuous improvements in raw data accuracy and new chemistries mean that less sequencing depth is needed for reliable results [37].

The present work can help to decide which adaptive sampling approach is best suited for analyzing specific clinical samples and questions. For example, when sequencing metagenomes with unknown microbial composition and host contamination, the depletion method is a better choice. On the other hand, the enrichment method might be helpful for metagenomics if only certain bacterial species are of interest as the higher rejection rates increase the sequencing depth further. Combining the real-time sequencing data stream of ONT with automated analysis pipelines, the turnaround time from sample collection to analysis and an appropriate treatment strategy can be reduced.

A few limitations must be noted. Our results were based on the library preparation with PCR amplification, for which we did not observe a noticeable bias when sequencing the microbial community standard. Still, these results might be different in other specimens. Due to the DNA isolation via bead beating and the PCR-based library preparation, we sequenced short DNA fragments of approximately 2500 bp only. Longer DNA fragments might improve the adaptive sampling’s decision-making, further increasing overall sequencing depth. Furthermore, the user should carefully select the provided reference genome(s) during adaptive sampling as, e.g., another human reference might slightly improve or worsen the overall depletion performance. Finally, raw read accuracy is currently at around 97.5% and might impact the read to reference mapping during sequencing and thus the adaptive sampling accuracy.

We strongly believe that adaptive sampling will prove exceptionally useful within clinical research and the individual microbiological and microbiological diagnostic approach in routine diagnostics. The increased information depth for compartment-specific human microbiomes in a physiologic and pathophysiologic context may change paradigms of antiinfective therapies in a personalized risk stratifying manner.

## 4 Methods

### Sample Selection

Vaginal swabs of adult pregnant women were admitted to the hospital with PPROM (hospitalization between 22+0 and 34+0 wks) and collected with sterile FLOQSwabs (Copan, Italy) within the PEONS trial (ClinicalTrials.gov NCT03819192) between September 2019 and March 2020. The institutional review board approved the study, and all participants signed a written informed consent. Swabs were immediately frozen at - 80°C until analysis was performed.

### DNA extraction

DNA was extracted from 75□Jμl ZymoBIOMICS Microbial Community Standard (Zymo Research Corporation, Irvine, CA, USA. Product D6300, Lot ZRC190633) using the ZymoBIOMICS DNA Miniprep extraction kit according to the manufacturer’s instructions. Similarly, the DNA from vaginal swabs was extracted using the ZymoBIOMICS DNA Miniprep extraction kit. The cell disruption was conducted for five minutes with the Speedmill Plus (Analytik Jena, Germany).

### Library preparation

DNA quantification steps were performed using the dsDNA HS assay for Qubit (Invitrogen, US). DNA of the microbial community standard was size-selected by cleaning up with 0.45× volume of Ampure XP buffer (Beckman Coulter, Brea, CA, USA) and eluted in 50□Jμl EB buffer (Qiagen, Hilden, Germany). The library was prepared from 1□Jμg input DNA using the SQK-LSK109 kit (Oxford Nanopore Technologies, Oxford, UK) and 5 ng using the SQK-RPB004 kit (Oxford Nanopore Technologies, Oxford, UK), according to the manufacturer’s protocol.

The sequencing library of clinical vaginal samples was prepared from 5 ng input DNA using the SQK-RPB004 kit (Oxford Nanopore Technologies, Oxford, UK), according to the manufacturer’s protocol.

### Nanopore sequencing

The microbial community standard was sequenced on the GridION using FLO-MIN106D Flow cells and the minknow-core-gridiron:4.1.2 software (all Oxford Nanopore Technologies). The Standard 48-hour script with active channel selection was applied.

Vaginal samples were sequenced by a standard 72-hour run script with and without adaptive sampling (depletion/enrichment). For ‘human depletion experiments’, the human reference genome GCA_000001405.28_GRCh38.p13 was used. A fasta reference file containing eight different bacteria species (*Aerococcus christensenii* (NZ_CP014159.1), *Fannyhessea vaginae* (NZ_UFSV01000001.1), *Gardnerella vaginalis* (NZ_PKJK01000001.1), *Lactobacillus iners* (NZ_AEKI01000028.1), *Mageeibacillus indolicus* (NC_013895.2), *Prevotella intermedia* (NZ_CP024727.1), *Prevotella jejuni* (NZ_CP023863.1), and *Ureaplasma parvum* (NC_010503.1)) was used for enrichment experiments. We performed a ‘Flow cell refuel’ step after approx. 18-20 hours of runtime by adding 70 μl of a 1:1 water-SQB buffer (Oxford Nanopore Technologies) to the flow cell SpotON port.

### Nanopore basecalling

Reads of the microbial community standard were basecalled using Guppy v4.2.2 GPU basecaller (Oxford Nanopore Technologies) during sequencing using the high accuracy basecalling model.

### Bioinformatics analysis

Sequencing data of the microbial community standard were mapped using minimap2 v.2.19 against the reference organisms’ sequences provided by Zymobiomics [22]. Subsequently, the base pairs sequenced per organism and per sequencing method were compared to each other.

The basecalled reads generated during adaptive sampling were extracted from the fastq files *via* samtools v1.19 and split based on the read_until.csv-file into three categories: ‘accepted’, ‘rejected’ and ‘no decision’. The reads were taxonomically classified *via* centrifuge v1.0.4 (updated centrifuge database h.v.b.a 01.2021 [27]).

## Author Contributions

Conceptualization,MM and CB.

Experiment conduction, MM

Figures created by MM and CB

JZ,JP,AV,ES,OM,MWP,RE

All authors actively participated in the writing and editing of the manuscript. All authors have read and agreed to the published version of the manuscript

## Acknowledgment

We sincerely thank the participating pregnant women for their support of this study. We also thank Yvonne Heimann and Dr. Kristin Dawcynszki from the PEONS study team for sampling and providing the swabs.

## Ethical approval and patient consent

This study was approved by ethical committees at University Hospital Jena (No. 2018-1183), University Hospital Halle/Saale (No. 2019-012), and University Hospital Rostock (No. A 2019-0055). Eligible women, who participated in the study, were informed about the study, applied procedures, and any risks due to sampling by a physician and gave their written consent. Participants were also informed that they could withdraw from the study at any time.

## Data availability

Centrifuge database available at: https://osf.io/5zv8t/.

Data on the patient microbiomes cannot be made publicly available for reasons of data protection, but can be exchanged bilaterally upon request and mutual non-disclosure agreement/cooperation agreement. For inquiries, please contact the corresponding author.

---

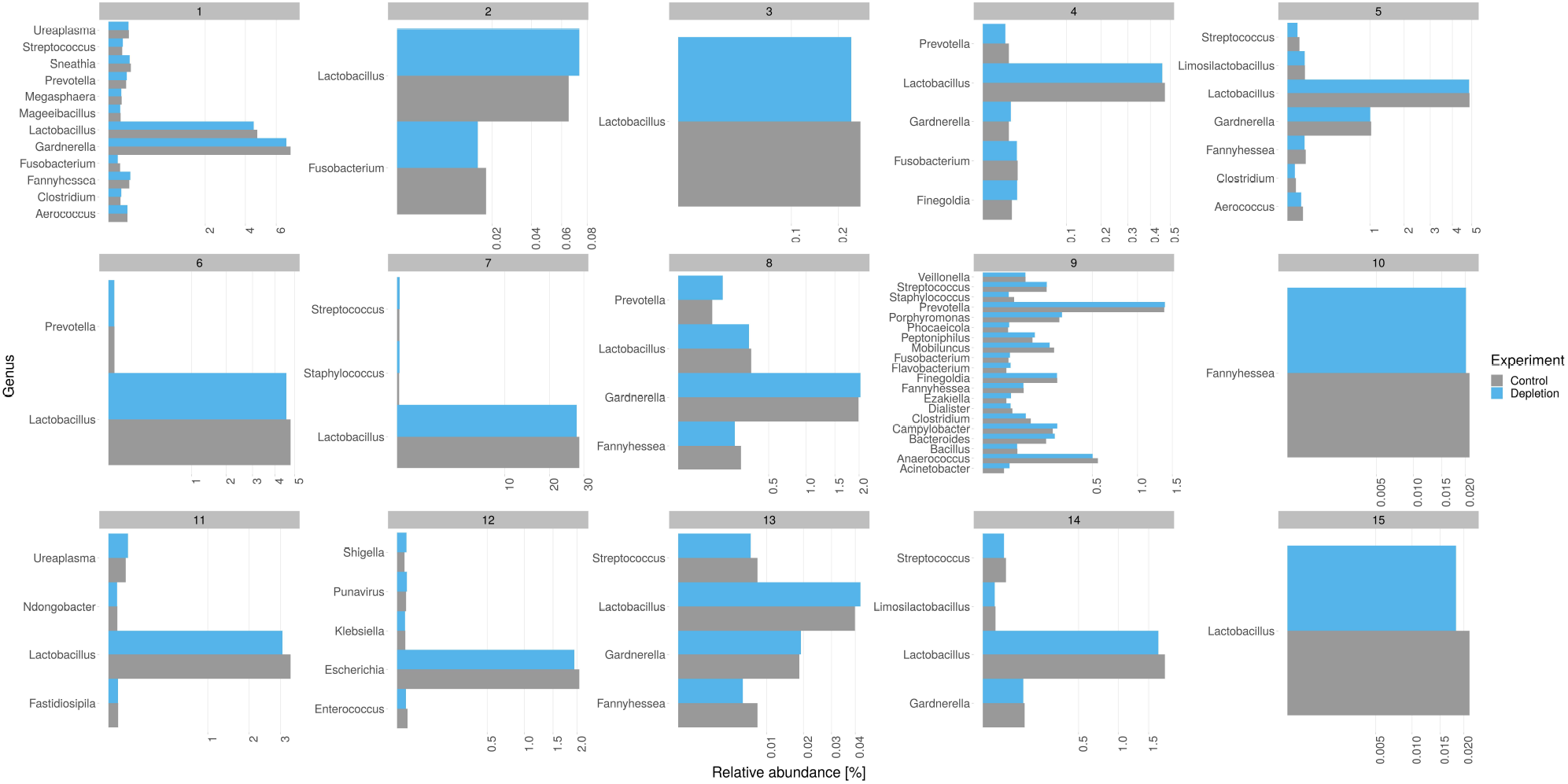

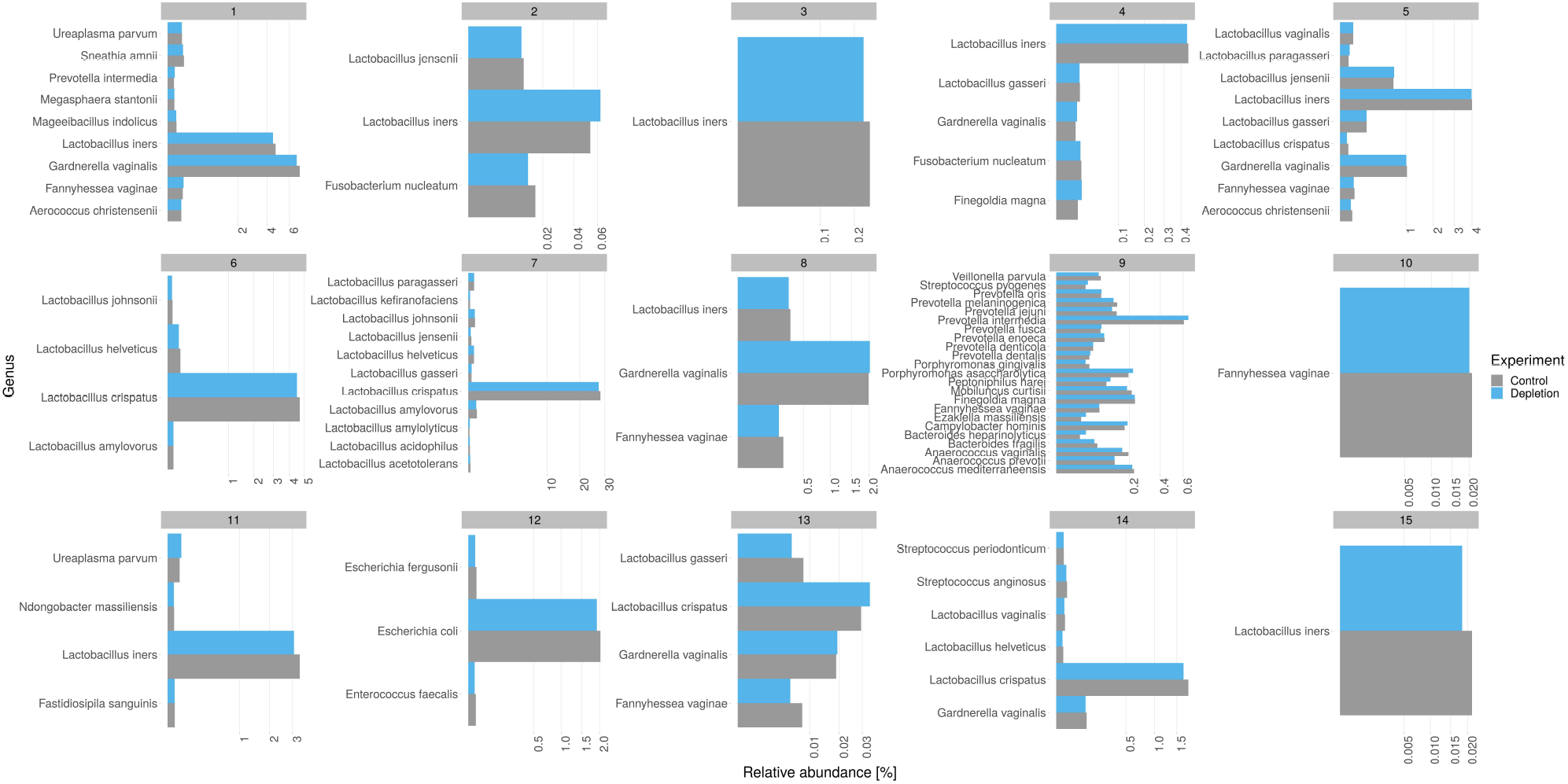

