## Supplementary Material S1 and S2 for "Evaluation of microbiome enrichment and host DNA depletion in human vaginal samples using Oxford Nanopore’s adaptive sequencing" (from another version of this preprint: version 1, 2021-09-15)

### 1 Supplementary Figures

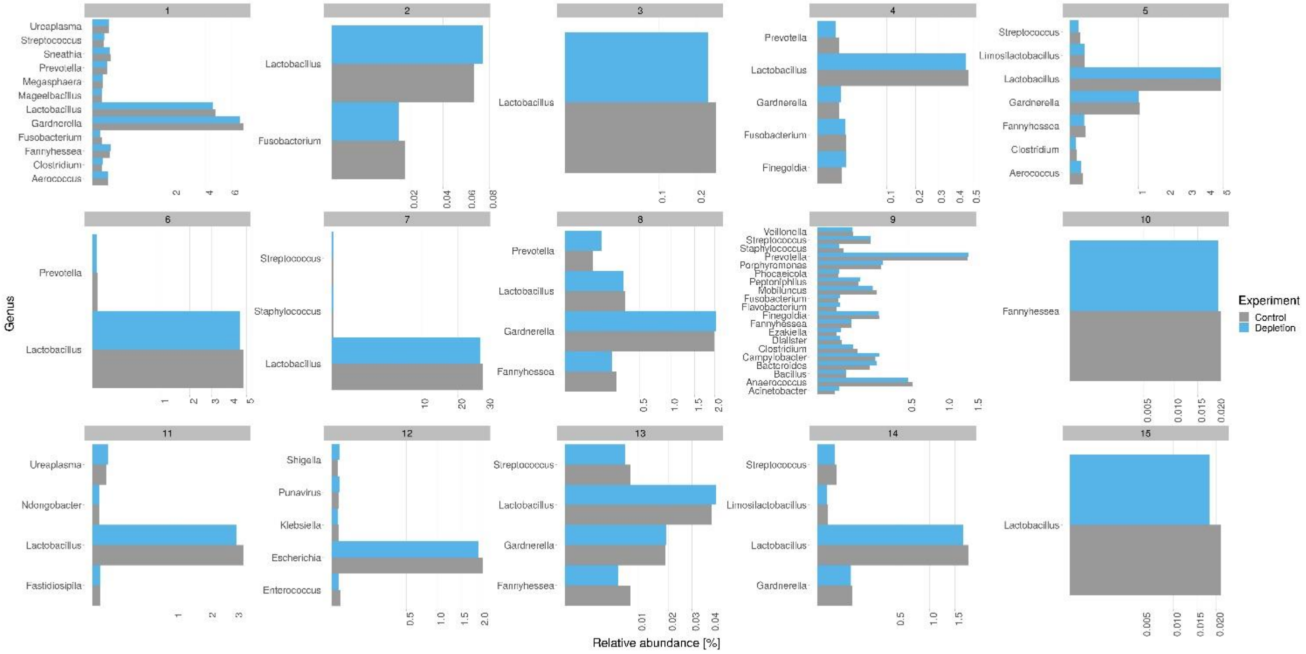


*Figure S1: Comparison of the relative abundance of bacterial genera for all vaginal samples (control in grey, depletion in blue).*

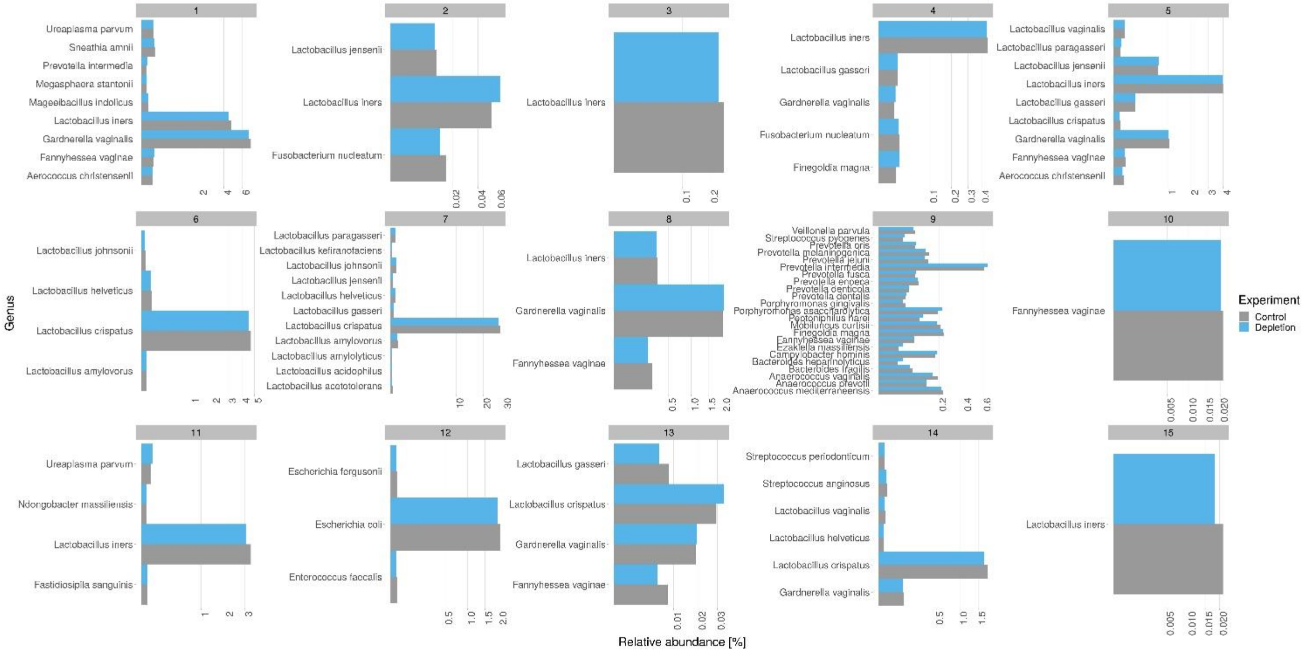


*Figure S2: Comparison of the relative abundance of bacterial species for all vaginal samples (control in grey, depletion in blue).*
